# Implications of variable synaptic weights for rate and temporal coding of cerebellar outputs

**DOI:** 10.1101/2023.05.25.542308

**Authors:** Shuting Wu, Asem Wardak, Mehak M. Khan, Christopher H. Chen, Wade G. Regehr

**Affiliations:** Department of Neurobiology, Harvard Medical School, Boston, MA 02115, USA

## Abstract

Purkinje cell (PC) synapses onto cerebellar nuclei (CbN) neurons convey signals from the cerebellar cortex to the rest of the brain. PCs are inhibitory neurons that spontaneously fire at high rates, and many uniform sized PC inputs are thought to converge onto each CbN neuron to suppress or eliminate firing. Leading theories maintain that PCs encode information using either a rate code, or by synchrony and precise timing. Individual PCs are thought to have limited influence on CbN neuron firing. Here, we find that single PC to CbN synapses are highly variable in size, and using dynamic clamp and modelling we reveal that this has important implications for PC-CbN transmission. Individual PC inputs regulate both the rate and timing of CbN firing. Large PC inputs strongly influence CbN firing rates and transiently eliminate CbN firing for several milliseconds. Remarkably, the refractory period of PCs leads to a brief elevation of CbN firing prior to suppression. Thus, PC-CbN synapses are suited to concurrently convey rate codes, and generate precisely-timed responses in CbN neurons. Variable input sizes also elevate the baseline firing rates of CbN neurons by increasing the variability of the inhibitory conductance. Although this reduces the relative influence of PC synchrony on the firing rate of CbN neurons, synchrony can still have important consequences, because synchronizing even two large inputs can significantly increase CbN neuron firing. These findings may be generalized to other brain regions with highly variable sized synapses.

## Introduction

The cerebellum is involved in behaviors ranging from balance, motor control and motor learning, to social and emotional behaviors (Hull and Regehr, 2022). Cerebellar disfunction has been linked to severe motor impairment and psychiatric disorders, including autism spectrum disorder, schizophrenia, bipolar disorder and depression (Argyropoulos et al., 2020; Schmahmann and Sherman, 1998). Within the cerebellar cortex, mossy fiber inputs from many brain regions excite granule cells that in turn activate Purkinje cells (PCs), which are the sole outputs. PCs project primarily to the cerebellar nuclei (CbN, also known as deep cerebellar nuclei or DCN), which then project to other brain regions. Clarifying how PCs control the firing of CbN neurons is a vital step in understanding cerebellar processing.

PCs are GABAergic and fire spontaneously at frequencies ranging from 10 spikes/s to over 100 spikes/s (Thach, 1968; Zhou et al., 2014). It was estimated that 40 PCs converge onto each CbN glutamatergic projection neuron to strongly suppress their firing (Person and Raman, 2012a). This is offset by the tendency of CbN neurons to fire spontaneously (Raman et al., 2000), and by numerous excitatory inputs from climbing fiber and mossy fiber collaterals (Najac and Raman, 2017; Wu and Raman, 2017). Excitatory inputs to CbN neurons are small and slow, and their number and firing rates are poorly constrained (Najac and Raman, 2017; Wu and Raman, 2017). Therefore, in this study we mainly focus on the control of CbN neuron firing by inhibitory inputs from PCs.

PCs are thought to convey information either with a rate code, or with a synchrony/timing code (De Zeeuw et al., 2011; Heck et al., 2013; Person and Raman, 2012a). There is considerable evidence that a rate code is used for some behaviors where the firing rates of PCs and CbN neurons are inversely correlated (De Zeeuw et al., 2011; Heck et al., 2013). However, for other behaviors, such inversely correlated firing is not observed, and a rate code does not apply (Armstrong and Edgley, 1984a, 1984b; McDevitt et al., 1987; Thach, 1970a, 1970b). This led to the hypothesis that synchronous firing of PCs that converge onto a CbN neuron could effectively regulate the rate and timing of CbN neuron firing (Gauck and Jaeger, 2000; Person and Raman, 2012a). Dynamic clamp studies of CbN neurons with 40 uniform sized PC inputs demonstrated the ability of PC synchrony to promote CbN neuron firing and precisely entrain spike timing (Person and Raman, 2012a). Whether PC outputs are encoded by firing rate, or by synchrony and timing, has been vigorously debated (Abbasi et al., 2017; Brown and Raman, 2018; Gauck and Jaeger, 2000; Heck et al., 2013; Herzfeld et al., 2023; Hoehne et al., 2020; Hong et al., 2016; Medina and Khodakhah, 2012; Payne et al., 2019; Person and Raman, 2012a, 2012b; Sedaghat-Nejad et al., 2022; Stahl et al., 2022; Sudhakar et al., 2015; Walter and Khodakhah, 2009; Wu and Raman, 2017).

Here we reexamine how PCs control the firing of CbN neurons. Previously it was thought that PC-CbN synapses are uniformly small, and that single PC inputs have little influence on the firing of CbN neurons. However, we find that unitary PC-CbN inputs have highly variable amplitudes, and some are very large. We used dynamic clamp and simulations to explore the implications of nonuniform input sizes. We find that for a given total inhibitory conductance, nonuniform input sizes led to highly variable inhibitory conductances and high basal CbN firing rates. Inputs of all sizes regulated the rate and timing of CbN neuron firing, but larger inputs had a bigger influence. Although PC synchrony had less influence on the firing rates of CbN neurons for nonuniform sized inputs because of higher basal firing rates, synchronizing several large inputs effectively regulated the CbN neuron firing. Thus, the nonuniform distribution of PC input sizes has important implications for how firing rates, timing and synchrony convey signals from the cerebellar cortex to the CbN.

## Results

### PC to CbN input sizes are highly variable in juvenile mice

To understand how PCs control the firing of CbN neurons, it is necessary to determine the distribution of the sizes of PC inputs. Previous studies found that PC to CbN inputs are relatively small and uniform in size, which could be approximated with a normal distribution. In order to determine whether this is the case in both young and juvenile age, we recorded the sizes of many individual inputs in young (p10-20, n=74) and juvenile (p23-32, n=83) animals (**Fig. 1**). We cut brain slices, stimulated PC axons with an extracellular electrode placed far from the CbN, and recorded the evoked IPSCs in CbN glutamatergic projection neurons with whole cell recordings. We adjusted the stimulus intensity to stochastically activate an individual PC axon and isolate a unitary input. This approach is shown for three inputs onto the same CbN neuron for a juvenile mouse (P27) (**Fig. 1a-c**). As shown in the individual trials for a small input, stimulation evoked short latency IPSCs (720 pA) in some trials but not others (**Fig. 1a**, *upper, grey*). The average of successes (**Fig. 1a,** *upper, black*) and failures (**Fig. 1a,** *upper, blue*) are shown. This is also apparent in a plot of the IPSC amplitudes for each trial (**Fig. 1a,** *lower*) showing that the IPSCs were evoked in a fraction of the trials. Similar experiments are shown for a medium-sized input (**Fig. 1b,** 1690 pA) and a large-sized input (**Fig. 1c,** 6280 pA). To better evaluate the variability in the input sizes, we plotted the distributions of unitary PC to CbN input conductances for young (**Fig. 1d**, P10-20) and juvenile (**Fig. 1e**, P23-32) mice. Input sizes were somewhat variable in young mice (**Fig. 1d**), but in juvenile mice there was considerably more variability, primarily because of the presence of many medium and large inputs (**Fig. 1e**). This is evident in a comparison of the cumulative histograms of the IPSC conductances for young and juvenile animals, which showed significantly different distributions (**Fig. 1f**; p<0.0001, Kolmogorov–Smirnov test). The distribution of input sizes we observed in P10-20 animals was similar to previous studies (P13-29, n=30) (Person and Raman, 2012a), and the skewed distribution of input sizes was only apparent when many individual inputs were characterized in P23-32 mice.

**Figure 1.**
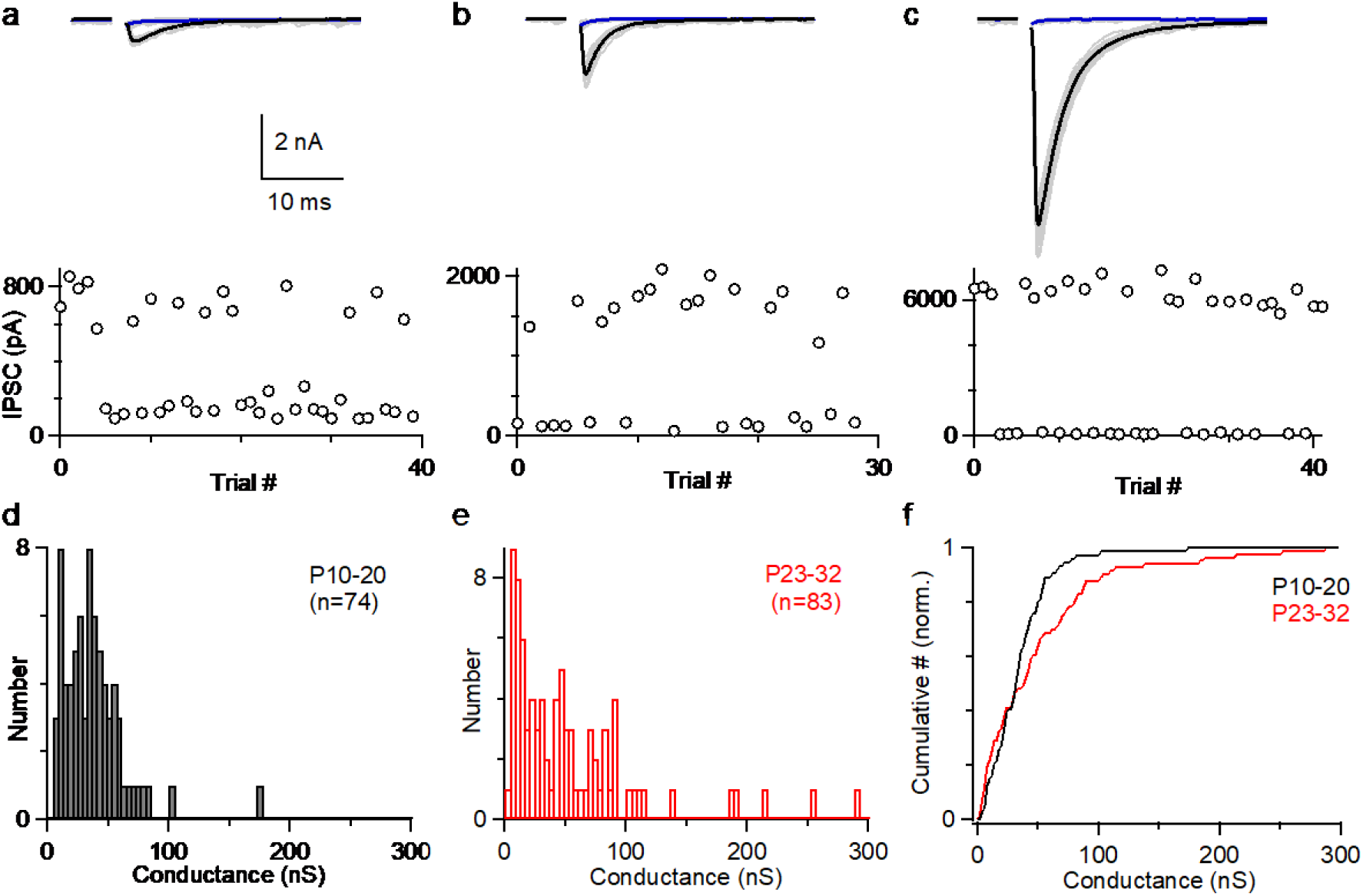
The amplitudes of Individual PC to CbN inputs are highly variable. **a.** Example of a small PC-CbN input (P27). (*top*) Responses evoked with the same stimulus intensity are superimposed for 40 trials (*grey*), and the average of successes (*black*) and failures (*blue*) are shown. (bottom) IPSCs amplitudes are plotted as a function of trial number. **b.** As in **a**, but for a medium sized input onto the same cell. **c.** As in **a**, but for a large sized input onto the same cell. **d.** Distribution of input sizes for PC-CbN IPSCs in young mice (P10-P20, n=74). **e.** Distribution of input sizes for PC-CbN IPSCs in juvenile mice (P10-P20, n=83). **f.** Normalized cumulative plot of conductances. The distributions of unitary input sizes in P10-20 animals and P23-32 animals were significantly different (p<0.0001) with a Kolmogorov–Smirnov test. The data Figures a-c were obtained for this study. The histogram of input sizes in (**d**, **e**) were based on new experiments (**d**, n=44; **e**, n=39), on (Turecek et al., 2017) (**e**, n=44), and (Khan and Regehr, 2020) (**d**, n=30).

### Using dynamic clamp and simulations to examine the effects of variable input sizes

The variability in the amplitudes of PC to CbN inputs in juvenile animals raised the issue of how the wide range of input sizes influences the way PCs control the firing of CbN neurons. We then designed dynamic clamp studies using highly variable PC-CbN input sizes. The PC-CbN unitary conductances we used in the dynamic clamp studies were based on our measurements with a few corrections. First, we used an internal with high concentration of Cl^-^ that provides superior voltage control to accurately determine the size of PC inputs. However, it has been shown in several studies that measurement of GABAR single-channel conductance or unitary conductance with high Cl^-^ based internals will result in a 2-3 times bigger value (Bormann et al., 1987; Gjoni et al., 2018; Sakmann et al., 1983). Therefore, we scaled down the unitary conductance amplitudes for physiological relevance (**Methods**). Second, we corrected the input sizes for the depression that occurs during physiological activation, which is approximately 40% of the initial size for a wide range of stimulus frequencies (Turecek et al., 2017, 2016). The excitatory conductances we used in the dynamic clamp studies were based on previous studies (Najac and Raman, 2017; Wu and Raman, 2017), with a relatively unconstrained frequency to pair with different inhibitory conductances (**Methods**).

The corrected distribution of input sizes used to guide our dynamic clamp studies is shown in **Fig. 2f** (*red*). For simplicity, instead of having a continuous range of input sizes, we approximated the distribution of PC input sizes in juvenile animals (**Fig. 1e, Methods**) with 16 small (3 nS), 10 medium (10 nS), and 2 large (30 nS) inputs (**Fig. 2f**, *grey*; total 200 nS, similar to the total conductances used in Person and Raman, 2012). The timing and frequency of the inputs were based on PC firing recorded in awake-behaving mice, with each input firing at approximately 80 Hz (**Fig. 2—figure supplement 1**, **Methods**). The resulting spike times were convolved with the corresponding unitary conductance size to generate the small (**Fig. 2b,** *green*), medium (**Fig. 2b,** *blue*), and large (**Fig. 2b,** *red*) conductances, which were then summed to generate the total inhibitory conductance arising from all inputs (**Fig. 2b,** *black*). The total inhibitory conductance showed large variations that were dominated by the medium and large inputs (**Fig. 2b**), that in turn produced large fluctuations in the membrane potential when injected into a CbN neuron (**Fig. 2c**). Spiking of the CbN neuron occurred mainly when the amplitude of the total inhibitory conductance was small (**Fig. 2b and 2c**).

**Figure 2.**
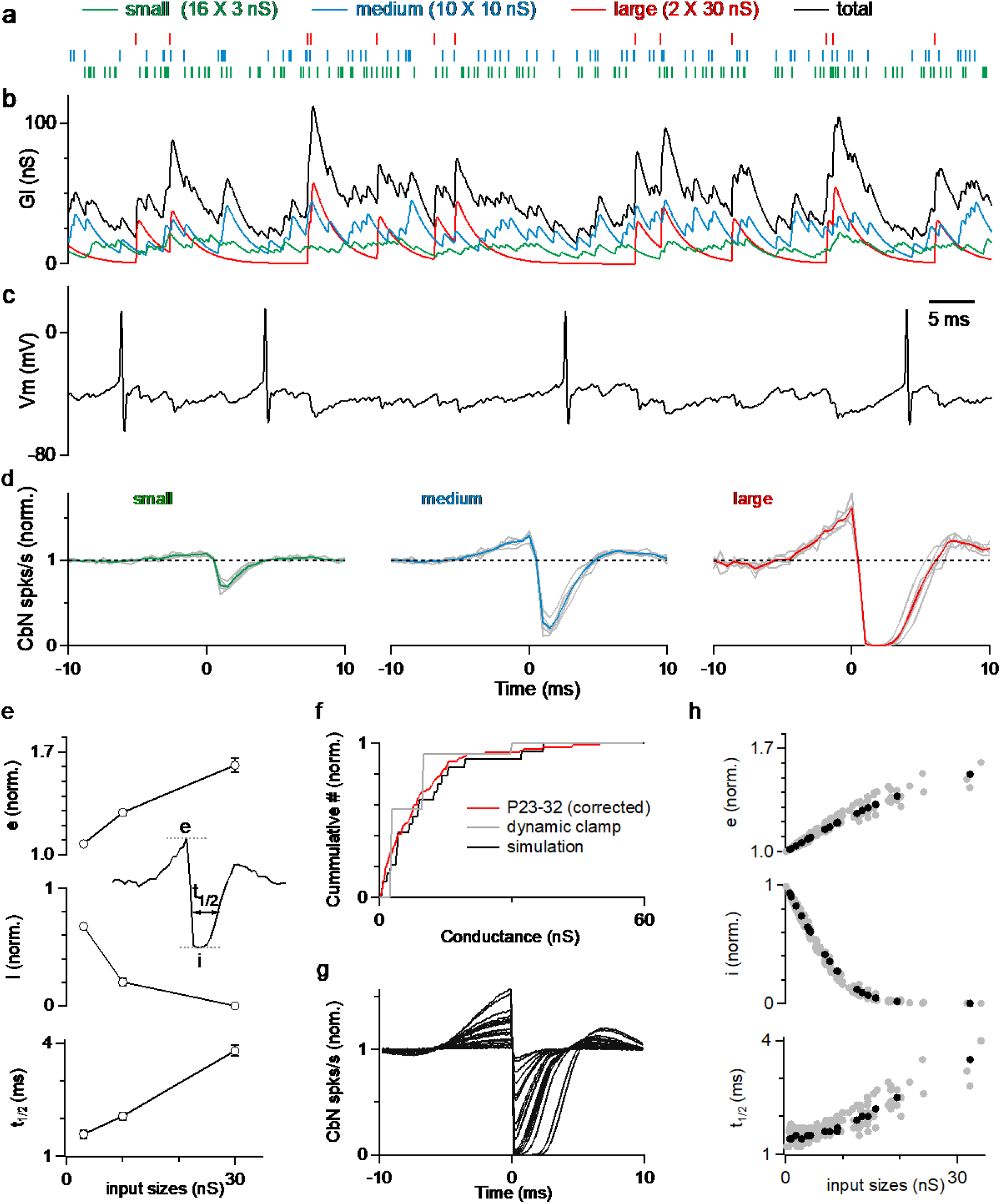
Large PC inputs powerfully influence the spike timing of CbN neuron firing. **a.** The corrected distributions of PC-CbN input sizes in juvenile animals (**f**, *red*) was approximated with 16 small inputs (3 nS, *green*), 10 medium inputs (10 nS, *blue*) and 2 large inputs (30 nS, *red*). Raster plots are shown for the spike times of the 16 small (green), 10 medium (blue), and 2 large (red) inputs used in dynamic clamp experiments. **b.** The total conductance waveform is shown (*black*) along with contributions from small (*green*), medium (*blue*) and large (*red*) inputs. **c.** Spikes in a CbN neuron evoked by the total conductance in (**b**) in dynamic clamp experiments **d.** The normalized cross-correlograms of input spiking and CbN neuron spiking for small, medium and large inputs for dynamic clamp experiments. Different cells (n=6, *grey*) are shown along with the average cross-correlograms (*colored traces*). **e.** Summary of the excitation (e), inhibition (i) and half-decay time (t_1/2_), as defined in the inset for the data in **d**. Inset shows how the parameters are determined. **f.** Cumulative histogram of all recorded inputs from Figure 1 for P23-32 mice corrected for depression and internal solution (*red*), the simplified input distribution used in dynamic clamp studies in **a** (*grey*), and inputs drawn from that distribution that were used in a simulation (*black*). **g.** Calculated cross-correlograms (normalized) for each of the different inputs used in a simulation. **h.** Summary of the excitation (e), inhibition (i) and half-decay time (t_1/2_), for a simulation as defined in the inset for the simulated cell in **g** (*black*), and for inputs to 9 other simulated cells with different distributions of inputs (*grey*).

**Figure 2—figure supplement 1.**
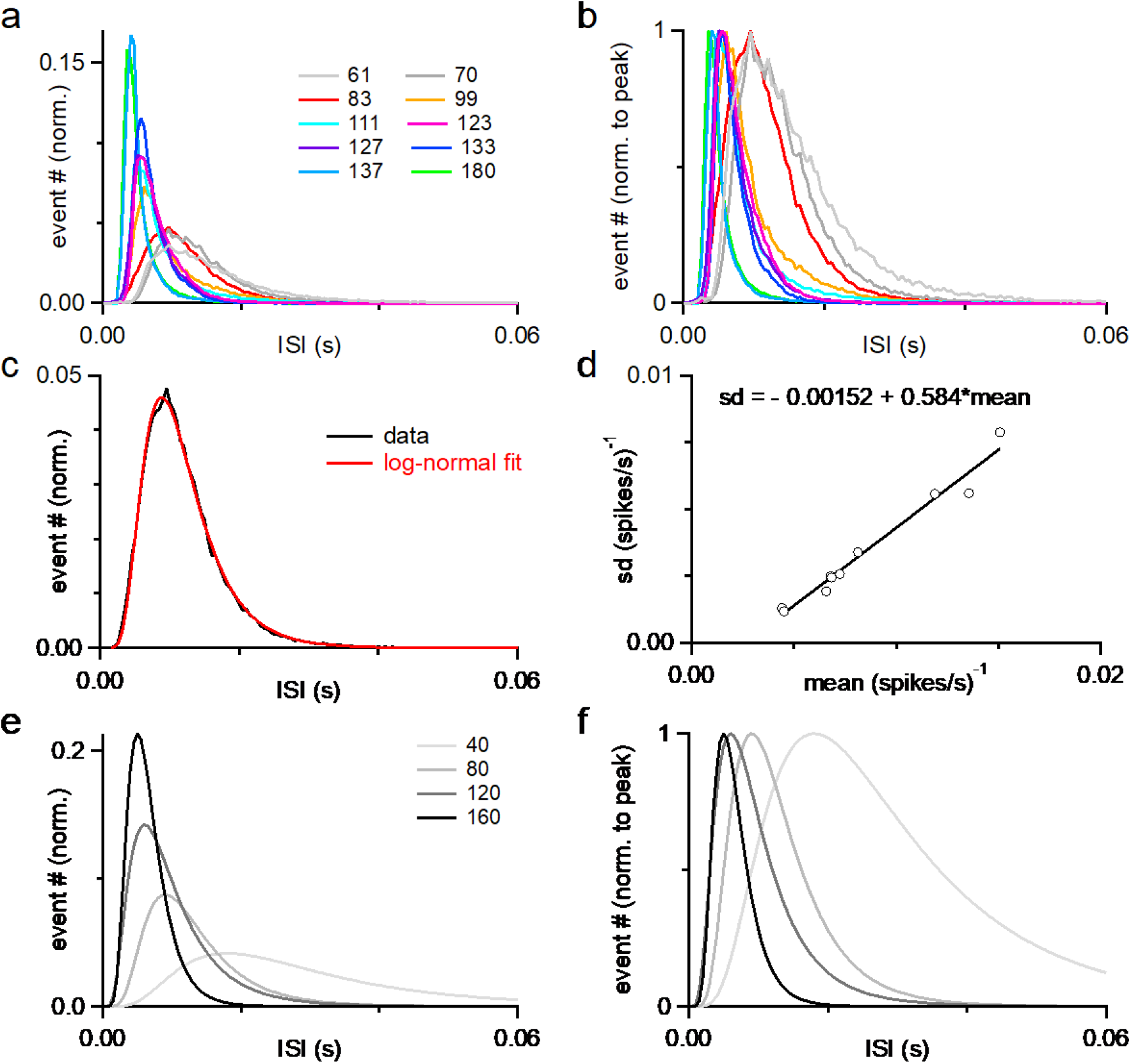
ISI histograms of PCs firing used in this study. **a.** Normalized ISI histograms of 10 PCs recorded *in vivo*, with their firing rates indicated in the legend. **b.** As in **a** but normalized to the peak. **c.** Example of a lognormal distribution fitting (red) to the ISI histogram of one PC recoded *in vivo* (black, 83 Hz). Similar fittings were performed for the other 9 PCs. **d.** The standard deviation (sd) as a function of the mean of the lognormal distribution fits to the 10 PCs was fitted as a linear function. Fits were performed in IGOR Pro to the function *exp*{−[*ln*(*x*⁄*x*_0_)/*width*]^2^}. The values of μ and σ for standard lognormal distribution are computed from *x_0_* and width using equations: 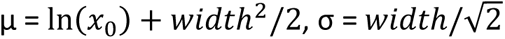. The values of mean and sd of the lognormal distribution fits were computed from μ and σ using equations: mean = *exp*(μ + σ ^2^/2) and 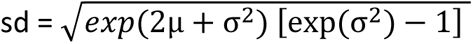. This linear function was used to determine the parameters for a PC firing ISI lognormal distribution with a desired firing rate. **e.** Four lognormal distributions representing artificial PC firing ISI distributions with different firing rates generated with the mean and sd from the linear function in **d**. **f.** As in **e** but normalized to the peak.

All inputs had effects on the timing of CbN neuron firing, but larger inputs had much bigger effects. As shown in the cross-correlograms of input timing and CbN neuron spiking, small inputs produced a small transient decrease in CbN neuron spiking (**Fig. 2d,** *left*), while medium sized inputs strongly reduced CbN neuron spiking (**Fig. 2d,** *middle*), and large inputs transiently shut down CbN neurons for approximately 2 ms (**Fig. 2d,** *right*). Intriguingly, the inhibition generated by inhibitory inputs was preceded by an increase in spikes that was particularly prominent for large inputs (**Fig. 2d**). The magnitude of excitation (**Fig. 2e**, *top*) and inhibition (**Fig. 2e**, *middle*), as well as the duration of inhibition (**Fig. 2e,** *bottom*) were all positively correlated with the size of the PC input. These findings highlight the ability of large PC inputs to control the timing of CbN neuron spiking.

We then performed simulations to extend these findings to a more realistic distribution of PC inputs sizes (**Methods**). We determined the sizes of PC input by assigning them values randomly drawn from the distribution of **Fig. 2f** (*red*) until the total inhibitory conductance reached 200 nS. An example distribution of PC input sizes used in a simulation is shown in **Fig. 2f**. The cross-correlograms of input timing and CbN neuron spiking for each of the PC inputs were determined, and they all showed a similar pattern, with an elevation followed by a suppression of CbN firing (**Fig. 2g**). The magnitudes of the excitation and inhibition of CbN firing had a similar dependence on the size of the PC inputs as shown for the example neuron (**Fig. 2h**, *black*), and for 9 other simulated neurons (**Fig. 2h**, *grey*). These simulations complemented our dynamic clamp studies and showed that the properties of cross-correlograms and their dependence on the size of the PC input can be extended to realistic distributions of input sizes.

### The autocorrelation of PC firing leads to disinhibition prior to inhibition of CbN neurons

The elevated spiking observed prior to the inhibition by a PC input is intriguing. If such a cross-correlogram was observed *in vivo*, it might be attributed to excitation of the CbN neuron that preceded PC inhibition, but that cannot be the case for our experimental conditions. Another possibility is that the refractory period of PC firing results in reduced inhibition and effective excitation. We tested this hypothesis by performing dynamic clamp experiments with the timing based on three different PCs recorded *in vivo*, and on an artificial Poisson distribution (**Fig. 3a**). As shown in the ISI histograms (**Fig. 3a**) and auto-correlograms (**Fig. 3b**), the three recorded PCs had refractory periods that were correlated with their firing frequency (∼3.5 ms for 49 Hz, ∼2 ms for 83 Hz, and ∼1.5 ms for 122 Hz firing), while the Poisson input did not have a refractory period. We then generated inhibitory conductances by convolving spike times with a 20 nS single input conductance. To compensate for the differences in input firing frequency, we adjusted the number of the inputs so that the total inhibitory conductances were approximately that same for all cases (12 x 20 nS at 49 Hz, 9 x 20 nS at 83 Hz, 6 x 20 nS at 122 Hz, and 9 x 20 nS for Poisson inputs). When input timing was based *on in vivo* recorded PCs that have a refractory period, the spike triggered average inhibitory conductances decreased prior to the elevation of the inhibitory conductance (**Fig. 3c**), and consequently the cross-correlograms showed elevated spiking in the CbN neuron (**Fig. 3d**). The speed and amplitude of this decrease in inhibitory conductance and the associated elevation in spiking depended on the PC firing statistics, with the faster firing inputs associated with larger, shorter-lived effects (**Fig. 3d**). The Poisson input that lacked a refractory period did not have a decrease in the inhibitory conductance, and therefore CbN firing was not elevated prior to inhibition in the cross-correlogram (**Fig. 3c and 3d,** *far right*). Thus, the autocorrelations of PC firing led to elevated CbN firing prior to suppression.

**Figure 3.**
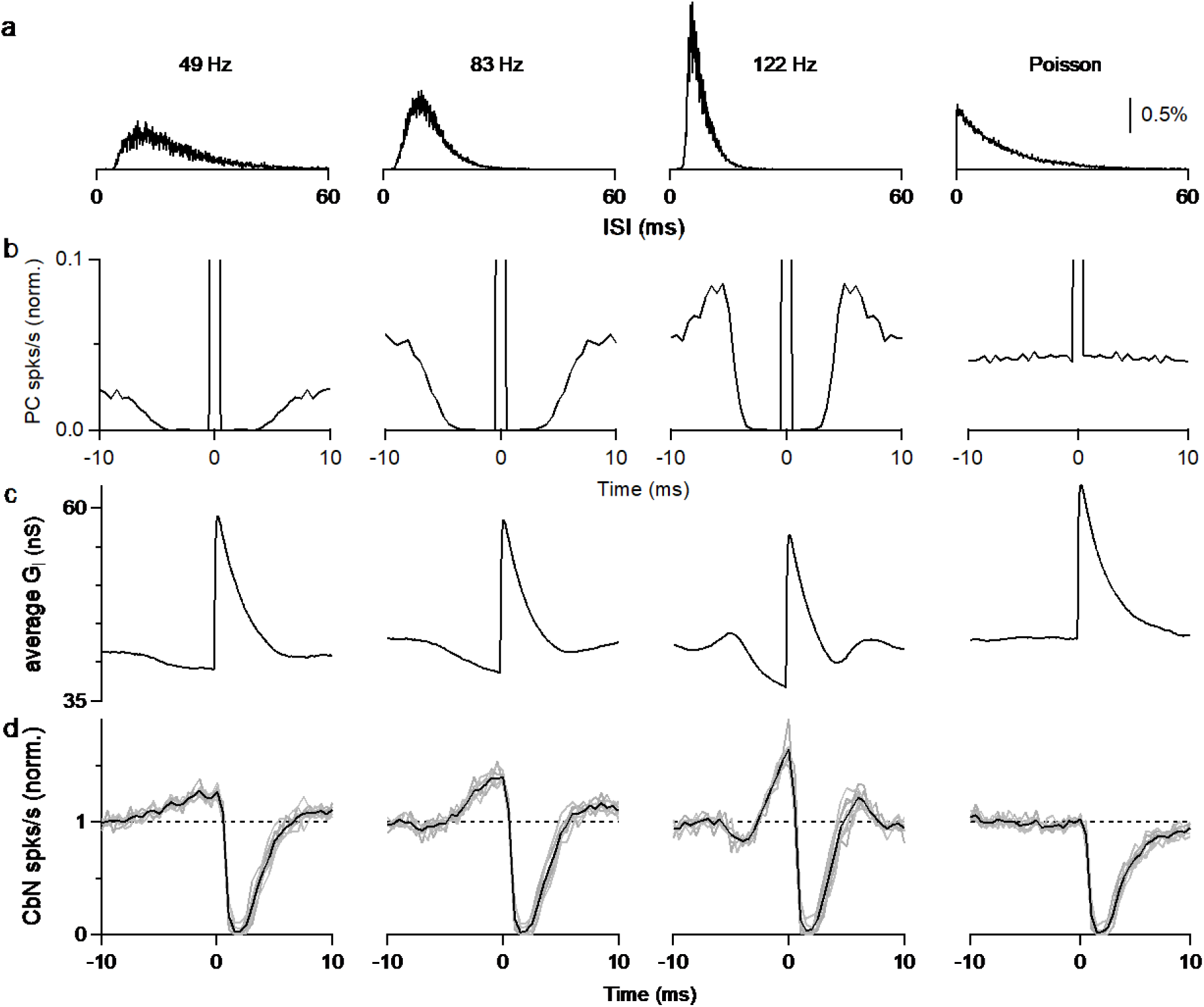
Autocorrelation of Purkinje cell firing leads to excitation prior to inhibition of CbN neurons. **a.** Interspike interval (ISI) histograms for three different PCs recorded *in vivo (left)*, and for an artificial Poisson input lacking a refractory period (*right*). **b.** Autocorrelation functions for the ISI distributions in **a**. At 0 ms, all graphs peak at 1 and graphs are truncated to allow better visualization. **c.** Calculated spike-triggered average inhibitory conductances for different cases (12 x 20 nS at 49 Hz, 9 x 20 nS at 83 Hz, 6 x 20 nS at 122 Hz, and 9 x 20 nS for Poisson inputs), with ISIs drawn from the corresponding distributions in **a**. **d.** Cross-correlograms are shown for PC inputs and CbN spiking for dynamic clamp experiments that used the distributions in **a**.

### The amplitude and coefficient of variation of PC inhibition regulate the firing rate of CbN neurons

Thus far we have shown the differential effects on CbN spike timing by different sized PC inputs. PCs also control the average firing rate of CbN neurons in addition to spike timing. We used dynamic clamp to determine how the amplitude and the coefficient of variation (CV) of the inhibitory conductance influence CbN firing. We began by examining how changing the amplitude of a constant inhibitory conductance altered the firing of CbN neurons (**Fig. 4a**). The firing rate of CbN neurons was approximately 200 spikes/s for an inhibitory conductance of 30 nS, but as the magnitude of the inhibitory conductance increased the spike rate decreased, and an inhibitory conductance of 65 nS silenced CbN firing (**Fig. 4a and 4b**). As expected, the CbN firing rate was inversely related to the amplitude of inhibitory conductance (**Fig. 4b**). We then examined how the variability in the inhibitory conductance influenced CbN spiking. We kept the total inhibitory conductance constant while varying the sizes and numbers of PC inputs that contribute to the total inhibitory conductance (**Fig. 4c**). We generated the total inhibitory conductance for the indicated numbers and sizes of PC synaptic inputs with the timing based on PC firing recorded *in vivo* (**Fig. 4c; Fig. 2—figure supplement 1**; **Methods**).

**Figure 4.**
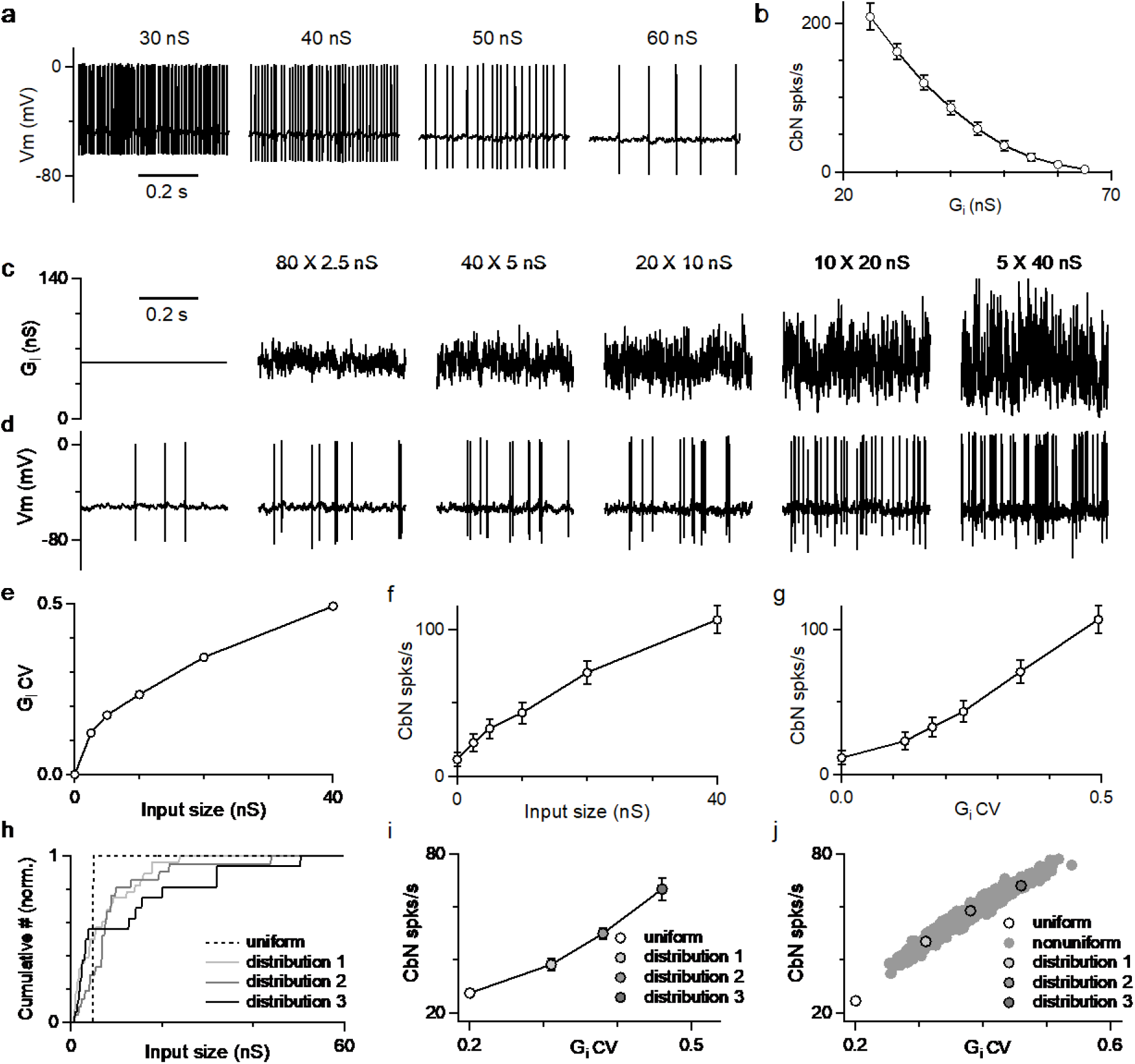
The amplitudes and fluctuations of the total inhibitory conductance both regulate the firing of CbN neurons. **a.** CbN neuron spiking observed for constant inhibitory conductances of the indicated amplitudes. **b.** Summary of CbN neuron firing rates (n=7 cells) during constant inhibitory conductances as in **a**. **c.** Inhibitory conductances with the same average conductance (56 nS) are shown for a constant conductance (*far left*), and for different cases with the numbers and sizes of inputs varied. **d.** CbN neuron spiking is shown for dynamic clamp experiments with the inhibitory conductances in **c**. **e.** The coefficient of variation (CV) of the inhibitory conductance is plotted as a function of the input size. **f.** CbN firing rate is plotted as a function of input size. **g.** CbN firing rate is plotted as a function of the CV of the inhibitory conductance. **h.** Normalized cumulative plots of conductances of three different input distributions (*solid lines*) drawn from the observed distribution (Fig. 2f, *red*), and for the 40 × 5 nS inputs (*dashed line*). **i.** CbN firing rates in dynamic clamp experiments that used the inhibitory conductances in **h** are plotted as a function of the CV of the inhibitory conductance. **j.** Simulated CbN firing rates based on 1000 different input distributions randomly drawn from the observed distribution of input sizes (*filled grey*) and for 40 × 5 nS inputs (*open circle*) are plotted as a function of the CV of the inhibitory conductance. The three distributions used in dynamic clamp experiments in **i** are highlighted.

While the average of the conductance wave was the same for all combinations of input sizes and numbers, as the size of the inputs increased, the conductance became increasingly variable (**Fig. 4c**), and the CV became larger (**Fig. 4e**). Evoked firing rates in CbN neurons were low for many small inputs, but as the number of inputs decreased and the size increased, the firing rate increased markedly (**Fig. 4d and 4f**). The firing rate was strongly dependent on the CV of the inhibitory conductance (**Fig. 4g**). These findings establish that increases in the magnitude of the average inhibitory conductance suppress firing, whereas increases in the variability of the inhibitory conductance promote firing.

The dependence of CbN firing on the variability of the inhibitory conductance prompted us to examine the influence of variable input sizes on the basal firing rate of CbN neurons. Three different distributions of PC input sizes drawn from the observed distribution of input sizes (**Fig. 2f**, *red*) with different CVs and a total conductance value of 200 nS (**Fig. 4h**, *solid lines*), together with 40 uniform 5 nS PC inputs (**Fig. 4h**, *dashed line*), were used to generate inhibitory conductances for dynamic clamp experiments. Even though the average conductance was the same, the firing rates for the three nonuniform input sizes were higher than for uniform inputs (38, 50, 67 vs. 27 spikes/s; **Fig. 4i**). The differences in the firing rates are readily explained by the CV of total conductances generated from different sized inputs (**Fig. 4i**). We also simulated the firing evoked by 1000 different distributions of PC input sizes drawn from the observed input size distribution with a total conductance size of 200 nS, and observed a broad range of firing rates that depended on the CV of the total inhibitory conductance (**Fig. 4j**). Both dynamic clamp experiments and simulations showed that for the same total inhibitory conductance, CbN neuron firing rates are always higher for nonuniform inputs than for uniform inputs as a result of the higher CV (**Fig. 4i and 4j**).

### Different sized inputs reliably transfer a rate code

The ability of different size PC inputs to convey a rate code to CbN neurons was unclear, given that both the magnitude and variability of total inhibition can influence the firing rates of CbN neurons. We addressed this issue using dynamic clamp experiments with small, medium, and large inputs (small: 16 × 3 nS = 48 nS; medium: 10 × 10 nS = 100 nS, and large: 2 × 30 nS = 60 nS), as in **Fig. 2**. We varied the firing rates of different sized inputs individually to determine how faithfully they convey a rate code. We began by varying the firing rates of all small PC inputs from 0 to 160 spikes/s while keeping the firing rates of medium and large inputs constant (**Fig. 5a,** *left;* **Fig. 2—figure supplement 1**, **Methods**). Elevating the firing rates of small inputs increased the amplitude (**Fig. 5b,** *left*) and decreased the CV (**Fig. 5c,** *left*) of the total inhibitory conductance, and suppressed the firing rate of CbN neurons (**Fig. 5d**, *left*). The CbN firing rate was inversely correlated with the firing rate of small inputs (**Fig. 5e**, *left*, *green*) and the inhibitory conductance amplitude (**Fig. 5e**, *middle*, *green*), and positively correlated with the CV of the inhibitory conductance (**Fig. 5e**, *right*, *green*). Varying the firing rate of all medium inputs (**Fig. 5a,** *middle*) had qualitatively similar effects on the conductance (**Fig. 5b,** *middle*), the CV (**Fig. 5c,** *middle*), and the CbN firing rate (**Fig. 5d,** *blue,* **Fig. 5e**, *blue*), but had larger effects because of their larger contribution to the inhibitory conductance. Varying the firing frequency of all large inputs (**Fig. 5a**, *right*) affected the total conductance (**Fig. 5b**, *right*) similarly to varying the small and medium inputs but had a very different influence on the CV (**Fig. 5c**, *right*). Eliminating the firing of either the small or medium inputs increased the CV of the inhibitory conductance, whereas eliminating the firing of the large inputs decreased the CV of the inhibitory conductance (**Fig. 5c**). This is consistent with large inputs being particularly effective at increasing the variability of the inhibitory conductance. However, it remained an open question whether the large inputs would be more effective at controlling CbN firing as a result of their greater influence on the variability and the CV of the inhibitory conductance (as in **Fig. 4g**), or if the amplitude of the inhibitory conductance amplitude (**Fig. 4b**) is the primary determinant of the CbN firing. We found the dependence of CbN firing rate on the amplitude of the total inhibitory conductance was highly consistent when the rates of either small, medium, or large inputs were varied (**Fig. 5e,** *middle*), but there were differences in the firing rate as a function of CV (**Fig. 5e,** *right*). These findings indicate that changes in the firing rate of PCs inputs are faithfully conveyed to the CbN primarily because of altering the total inhibitory conductance. Finally, to determine the influence of individual inputs on CbN firing, we calculated the slope of CbN output firing rate vs. the PC input firing rate for different size inputs and divided by the number of inputs (**Fig. 5f**). Compared to small and medium inputs, large individual PC inputs were much more effective at decreasing the firing rates of CbN neurons (**Fig. 5f)**. Therefore, the ability of a PC input to regulate CbN firing simply depends on its size.

**Figure 5.**
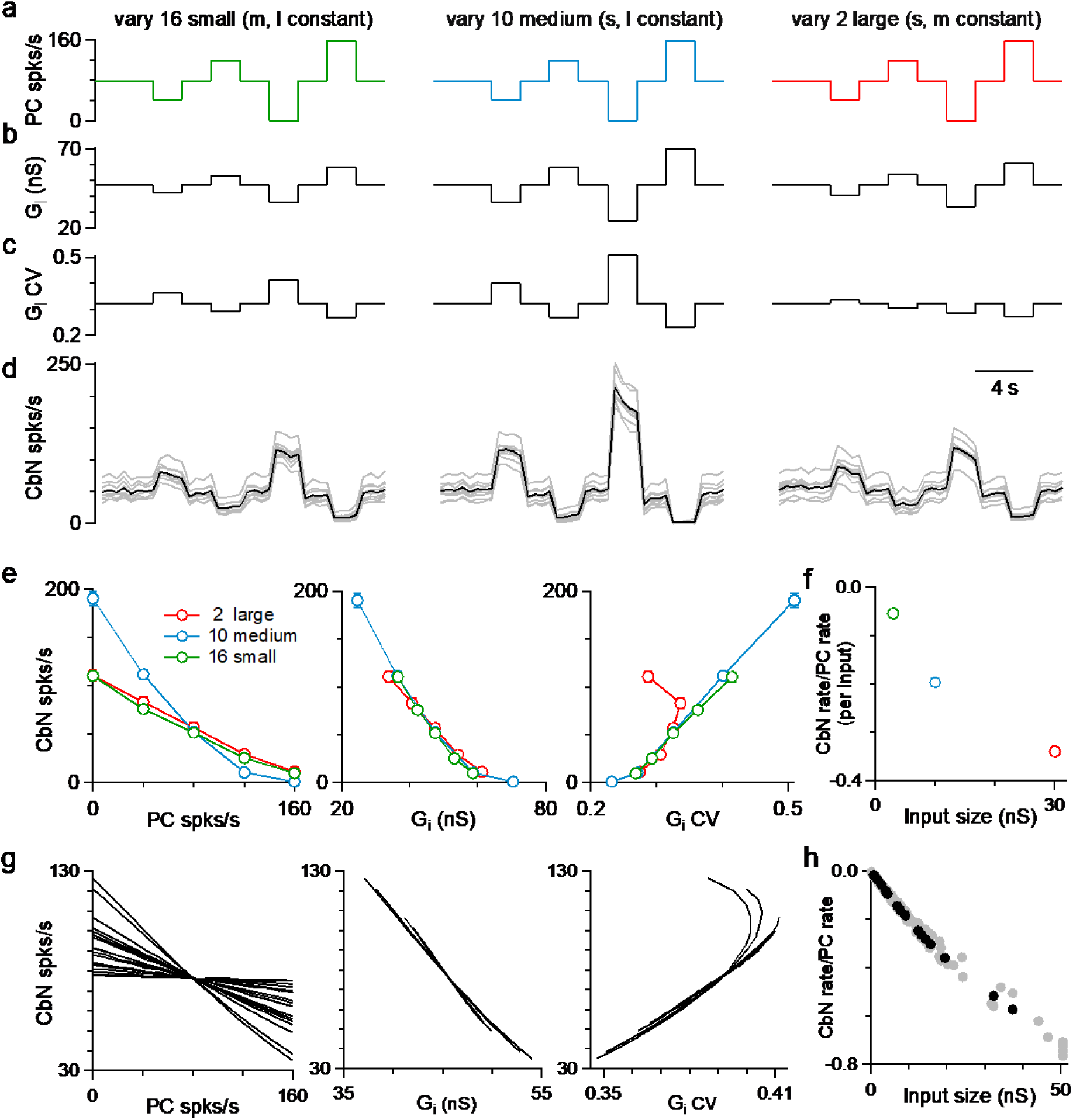
PC inputs effectively convey a simple rate code regardless of input size. Dynamic clamp experiments were performed with small (16 × 3 nS, green), medium (10 × 10 nS, blue) and large (2 × 30 nS, red) inputs. **a.** The firing rates of either small (left), medium, (middle) or large (right) inputs were varied every 2 s while the firing rates of the other inputs were maintained at 80 spikes/s. **b.** The resulting average inhibitory conductances for the three different conditions are shown. **c.** The resulting average CV of the inhibitory conductances are shown. **d.** The firing rates of CbN neurons with the conductances in **a-c** are shown as individual cells (n = 9, *grey*) and their average (black). **e.** The average firing rates of CbN neurons are plotted as a function of the rate of the PC inputs (left), the total inhibitory conductance (G_i_, *middle*), and the CV of the conductance (right), where the firing rates of either the small, medium or large inputs were changed. **f.** The slope of CbN output firing rate vs. PC input firing rate divided by the number of inputs are plotted for different size inputs. **g.** As in **e** but for simulations where the firing rates of each input were varied. **h.** As in **f** but for simulations for one CbN neuron (black) and other 9 CbN neurons (grey).

Simulations allowed us to examine the influences of individual PC inputs with a full range of sizes on the firing rates of CbN neurons. Simulations were similar to those of **Fig. 2g-h**, but with the firing rates for each input varied between 0 and 160 spikes/s while maintaining the average firing rates of other inputs at 80 Hz (**Methods**). Varying the firing rate of individual PC inputs led to an approximately linear, negatively correlated change in CbN firing rate (**Fig. 5g**, *left*). Single large inputs were surprisingly effective at regulating the firing rates of CbN neurons. In this example, varying a single 37 nS input from 0 to 160 spikes/s decreased the output firing frequency from 126 spikes/s to 35 spikes/s (**Fig. 5g**, *left*). Similar to the dynamic clamp experiments, there was a remarkably consistent relationship between the firing rates of CbN neurons and the total PC input conductances (**Fig. 5g**, *middle*), which was not observed for the CbN firing rates vs CV of the total inhibitory conductance (**Fig. 5g,** *right*). Lastly, the slope of the CbN output firing rate/ PC input firing rate was negatively correlated with the input size for the example neurons (**Fig. 5h**, *black*) and for 9 other neurons (**Fig. 5h**, *grey*). These simulations, along with the dynamic clamp studies, establish that individual PC inputs can regulate the firing rate of the CbN neuron, and that different sized inputs convey a rate code that depends on the amplitude of total inhibition.

### Synchrony of uniform or variable sized PC inputs

Previous dynamic clamp studies based on uniform PC input sizes revealed the potential importance of synchrony in regulating the firing of CbN neurons (Gauck and Jaeger, 2000; Person and Raman, 2012a). We extended this approach to assess the effects of synchrony for different sized PC inputs. We performed dynamic clamp experiments with both uniform sized inputs (**Fig. 6a**) and variable sized inputs (**Fig. 6b**) in each cell. We first looked at the effects of synchrony on the firing rate of CbN neurons. In the absence of synchrony, uniform sized inputs generated a baseline conductance with low variability and CV (**Fig. 6a***ii* **and 6a***iii*) that resulted in a low baseline firing rate of CbN neurons (25 spikes/s; **Fig. 6a***iv*). Synchronizing either 25% or 50% of the uniform inputs (**Fig. 6a***i*) increased the variability (**Fig. 6a***ii*) and the CV (**Fig. 6a***iii*) of the total inhibitory conductance, which then resulted in a robust increase in the firing rate of CbN neurons (**Fig. 6a***iv*). The CbN firing rate increased from the baseline firing rate of 25 spikes/s to 45 spikes/s with 25% synchrony, and 88 spikes/s with 50% synchrony (**Fig. 6a***iv***; Fig. 6c,** *black*). The increases in CbN firing rate were 1.8-fold higher for 25% synchrony and 3.5-fold higher for 50% synchrony, respectively (**Fig. 6a***v*).

**figure 6.**
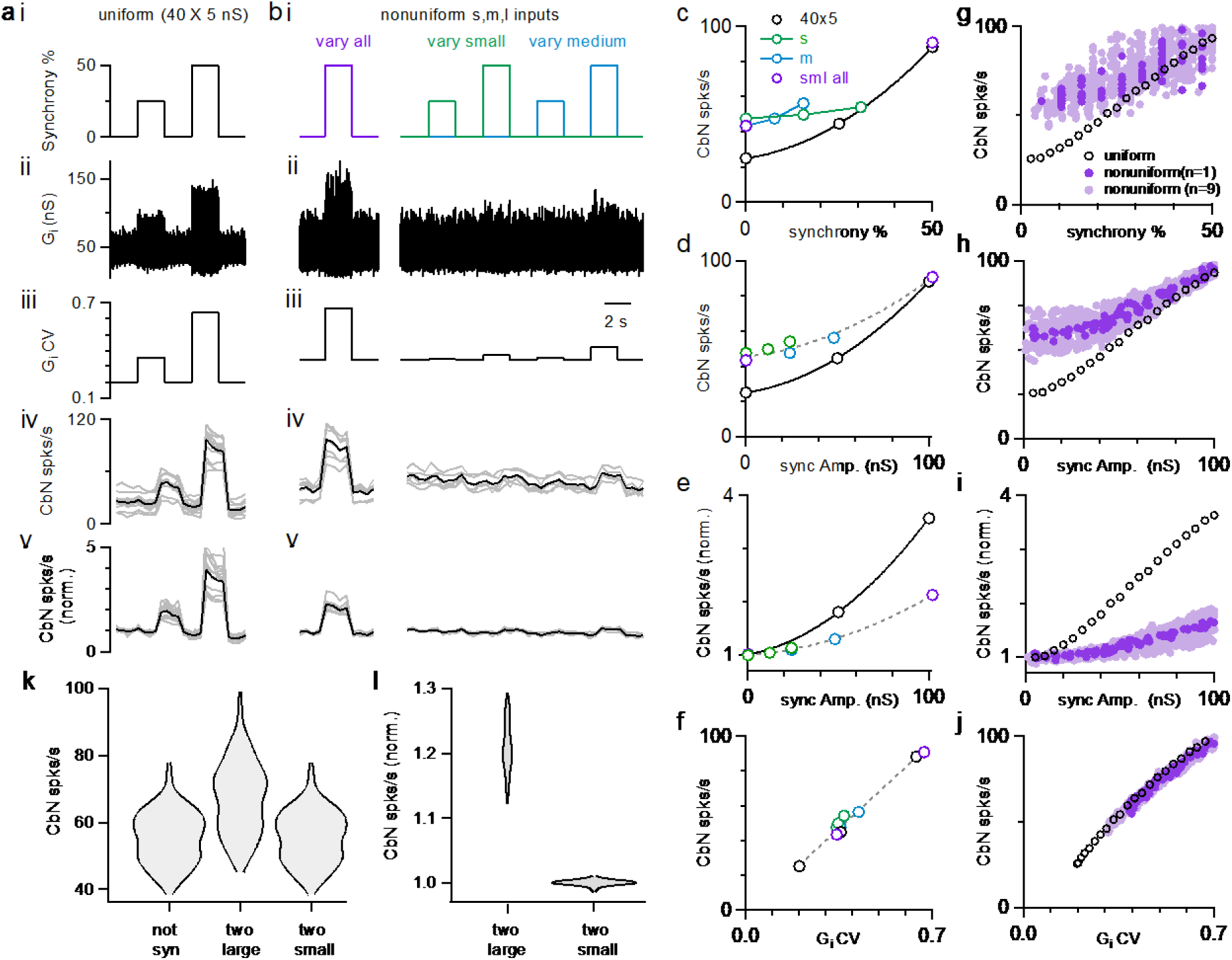
The influence of PC synchrony on the firing rates of CbN neurons for uniform and nonuniform sized PC inputs. Dynamic clamp experiments were conducted with either uniform (**a**) or nonuniform inputs (**b**) in which the conductance was kept constant, but the synchrony of the inputs was varied. **a.** i. The extent of synchrony of uniform sized inputs (40 × 5 nS) is varied. ii. The inhibitory conductance in which the synchrony of inputs is varied as in **a**. iii. The CV of the inhibitory conductance. iv. CbN neuron firing rate with the conductance (average, *black*; individual cells, n=14, *grey*). v. The normalized firing rate for the cells in **iv**. **b.** Similar experiments were performed on the same cells as in **a**, but for inputs of nonuniform sizes (small: 16 × 3 nS, *green*; medium: 12 × 8 nS, *blue*; large: 2 × 30 nS, *red*). In some cases, all types of inputs were synchronized (*purple*), in others only the small inputs (*green*) or the medium inputs (*blue*) were synchronized. **c.** CbN neuron firing rate as a function of the percentage of synchronous inputs. **d.** CbN neuron firing rate as a function of the total amplitude of synchronous inputs. **e.** The normalized CbN firing rates as a function of the amplitude of the synchronized inputs. **f.** CbN neuron firing rate as a function of the CV of the inhibitory conductances. **g. -j**. Similar plots to **c-f**, but for simulations with different sized inputs (dark purple, based on a single distribution, light purple based on 9 other different distributions) and uniform inputs (40 × 5 nS, black circles). **k.** Violin plots showing the simulated CbN neuron firing rate with 100 different distributions of different sized inputs (not syn), and the two largest inputs or the two smallest inputs synchronized. **l.** As in **k** but normalized to the not synchronized firing rate.

For nonuniform inputs (small: 16 × 3 nS; medium: 8 × 12 nS; large: 2 × 30 nS), the generated conductance was more variable (**Fig. 6b***ii* **and 6b***iii*), resulting in a higher baseline firing rate of CbN neurons (43 spikes/s; **Fig. 6b***iv*)), which is similar to **Fig. 4h-i**. Synchronizing 50% of all inputs (small, medium, and large inputs, **Fig. 6b***i*, *left*), increased the variability (**Fig. 6b***ii*, *left*) and CV (**Fig. 6b***iii, left*) of the conductance, which then elevated the firing of CbN neurons from 43 spikes/s to 91 spikes/s (**Fig. 6b***iv, left*; **Fig. 6c**). The firing rates for 50% synchrony with nonuniform and uniform inputs were comparable (**Fig. 6a***iv* **and Fig. 6b***iv*), but because baseline firing rates were higher for nonuniform inputs, their relative increase in CbN firing was much lower (2.1-fold for nonuniform inputs compared to 3.5-fold for uniform inputs, p<0.0001; **Fig. 6a***v* **and Fig. 6b***v*). We also examined the effect of synchronizing 25% or 50% of small or medium inputs (**Fig. 6b***i*, *right*). Synchronizing 25% or 50% of either small or medium inputs barely increased the variability and CV of the conductance, and led to minimal increases in the firing of CbN neurons (**Fig. 6b***ii-v*, *right*). These observations suggest that the effects of PC synchrony primarily depend on the total amplitude of synchronized inputs (**Fig. 6d and 6e**). The firing rates of nonsynchronized and synchronized inputs, for either uniform or nonuniform input sizes are readily explained by the influence of the CV of the inhibitory conductance on the CbN firing rate (**Fig. 6f**).

Simulations allowed us to explore the effects of a full range of synchrony levels, which would be impractical to test experimentally (**Fig. 6g-j**). We examined uniform inputs (40 × 5 nS) and determined the effect of synchronizing 1, 2, …20 inputs (**Fig. 6g-j,** *black open circles*). We also examined nonuniform sized inputs drawn from the full distribution of sizes (**Fig. 2f,** *red*), with a total conductance of 200 nS, and determined the firing rates evoked when we synchronized different combinations of inputs (**Fig. 6g-j,** *dark purple:* for one distribution of input sizes*; light purple:* for 9 other distributions of input sizes). The effects of synchrony on firing rates, and the dependence on the CV of the conductance were qualitatively similar for simulations (**Fig. 6g-j**) and dynamic clamp experiments (**Fig. 6c-f**). There was a great deal of scatter in the firing rates plotted as a function of the percentage of synchronized inputs (**Fig. 6g**, *dark purple*), but much less scatter when the firing rates were plotted as a function of the total amplitude of the synchronized inputs (**Fig. 6h**, *dark purple*). This is consistent with our observation that synchronizing many small or medium size inputs does not have much effect on the firing frequency of CbN neurons (**Fig. 6b***i-v, right;* **Fig. 6c-d**). For nonuniform inputs, synchrony has less influence on firing rates, because the baseline firing rates are higher (**Fig. 6e and 6i**). Nonetheless, synchrony of even a small number of large inputs can robustly elevate firing rates. As shown in simulations, synchronizing the two largest inputs can elevate CbN firing rate by approximately 20%, while synchronizing the two smallest inputs barely changed CbN firing rate (**Fig. 6k and 6l**).

We also examined the influence of PC synchrony on the spike timing of CbN neurons in dynamic clamp experiments of **Fig. 6**. Synchrony strongly altered the cross-correlograms of PC inputs and CbN firing, especially for the uniform sized inputs (**Fig. 7a and 7b**). For uniform inputs (5 nS), single unsynchronized inputs weakly influenced the spike timing of CbN neurons (**Fig. 7a**, *left*). 50% synchrony of uniform inputs silenced CbN neurons for several milliseconds, and this was proceeded by a prominent increase in CbN spiking (**Fig. 7a**, *right*). For asynchronous nonuniform inputs, as in **Fig. 2**, the influence of individual inputs on the timing of CbN firing depended on the size of the input, and single large inputs could have a very large influence on the timing of CbN spiking (**Fig. 7b**, *left; red*). Synchronizing 50% of all nonuniform inputs generated a cross-correlogram similar to that of synchronizing 50% uniform inputs (**Fig. 7a and 7b**, *right*). The strength of excitation (**Fig. 7c,** *top*), the amplitude of inhibition (**Fig. 7c,** *bottom*), and the duration of inhibition (**Fig. 7d**) were dependent on the synchrony amplitude, as was the case for simulations (**Fig. 7e and 7f**). For both dynamic clamp experiments and simulations, synchronizing the same amplitude of inputs led to larger timing effects for uniform inputs than for nonuniform inputs (**Fig. 7c-f**). This is likely because the large inputs present in the nonuniform case are also effective at controlling spike timing. These findings establish that for nonuniform inputs, the effects of synchrony on CbN spike timing depends on the total size of synchronized inputs.

**Figure 7.**
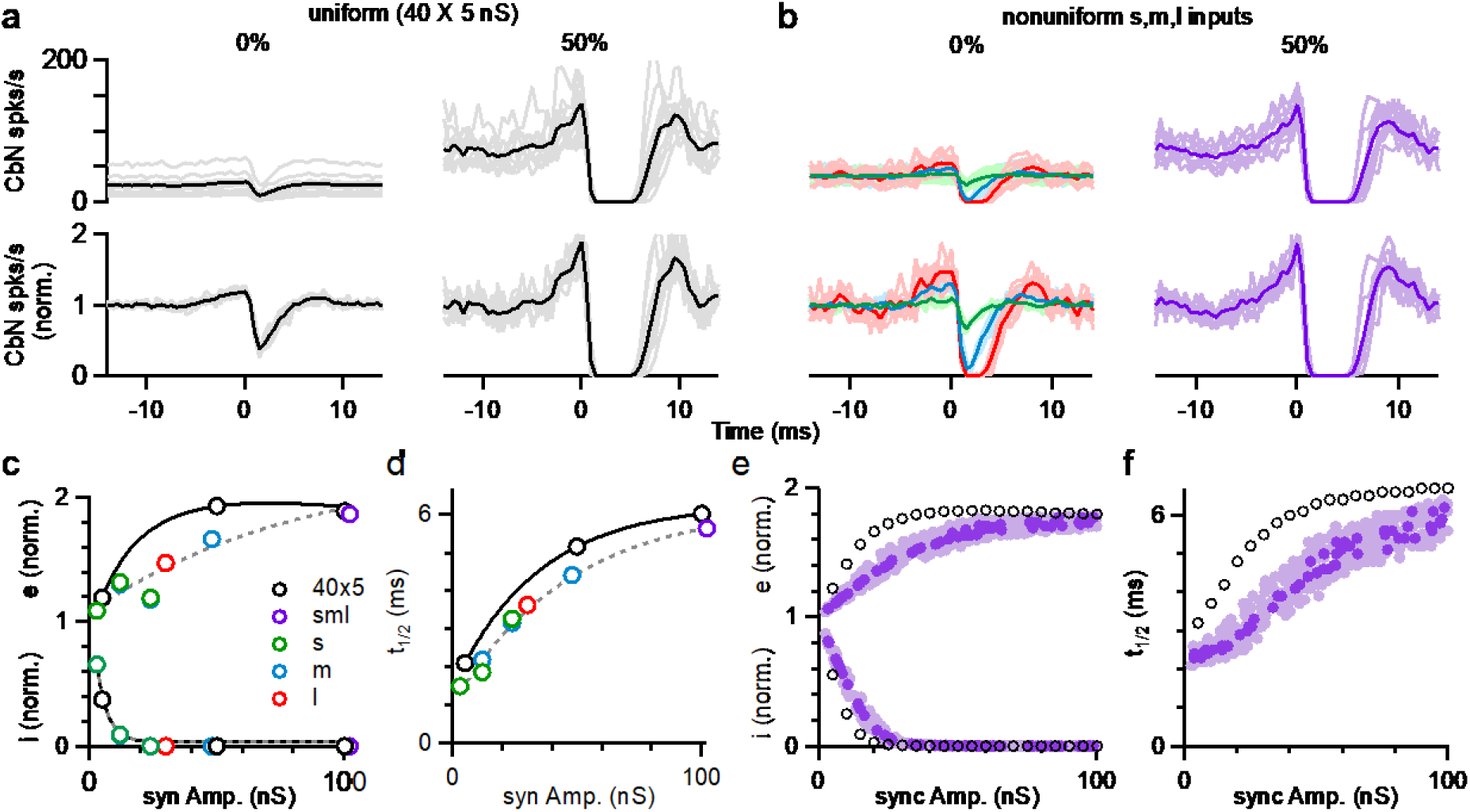
The influence of PC synchrony on the spike timing of CbN neurons for uniform and nonuniform sized PC inputs. **k.** *(top*) The cross-correlograms of PC input and CbN neuron spiking for nonsynchronous inputs (*left*) and 50% synchronous inputs (*right*) with uniform sized inputs. Individual cells *(grey*) and averages (*black*) are shown. (*lower*) As above, but normalized to the baseline firing rate. **l.** As in **a** but for different sized inputs (as in Fig. 6). The cross-correlograms of small (*green*), medium (*blue*), and large (*red*) inputs for unsynchronized inputs (*left*) and 50% synchronous inputs (*right*, *purple*) are shown. **m.** Summary plots of the excitation (e), inhibition (I) as a function of the amplitude of synchronized inputs. **n.** As in **c** but for half-decay time (t_1/2_). **o.** -**f**. As in **c-d**, but for simulations.

## Discussion

Our finding that single PC to CbN inputs are highly variable in size has important implications for how PCs control the firing of CbN glutamatergic projection neurons. We find that PC to CbN synapses are well suited to both implement rate codes and precisely regulate the timing of CbN neuron firing. Nonuniform input sizes increase the variability of the inhibitory conductance, which in turn elevates the basal firing rates of CbN neurons. Therefore, the relative influence of synchrony on CbN neuron firing rates is smaller for nonuniform inputs than for uniform inputs, but synchronizing even a small number of large inputs can significantly increase CbN neuron firing.

### Comparison to previous studies

The approach we took to examine the influence of PCs on CbN neuron firing is similar to that used in previous studies of the influence of PCs on CbN neuron firing (Person and Raman, 2012a), but we build upon those studies in several ways. As in earlier studies, we determined the properties of individual PC inputs and confirmed that they had very rapid kinetics (Person and Raman, 2012a). We then used dynamic clamp to study the firing of CbN neurons using measured PC-DCN conductances in combination with excitatory conductance (Wu and Raman, 2017). We confirmed that many small desynchronized PC inputs are highly effective at suppressing CbN firing, and that synchrony effectively controls the timing and rate of PC firing for small uniform sized PC inputs (Person and Raman, 2012a). Our observation that in somewhat older animals (P23-32) there is a skewed distribution of input sizes, and that some inputs are very large, had important implications. A single large input is functionally equivalent to many small, perfectly synchronized inputs. This allows PC inputs to influence the rate and timing of CbN firing as previously proposed (Person and Raman, 2012a), but without requiring a high degree of PC synchrony. We also extended previous studies by using realistic PC firing patterns in our dynamic clamp experiments. This allowed us to determine cross correlograms for individual inputs, and led to the finding that individual inputs have a surprisingly large effect on CbN firing. Previously, it was shown that synchronous PC inputs initially suppress CbN neuron firing, which is *subsequently* followed by an increase in firing (Person and Raman, 2012a). Here we find that for spontaneously firing PCs, a period of strong disinhibition *precedes* suppression by a PC input. Lastly, we complemented our dynamic clamp studies with simple simulations that faithfully reproducing our experimental findings, which allowed us to explore the influence of different parameters.

### The variable distribution of PC-CbN input sizes

We showed that there are developmental differences in the distributions of PC-CbN synapse sizes. In young animals (P10-P20, **Fig. 1d and 1f**, n=74), unitary PC inputs are relatively small and uniform in size, whereas in P23-32 mice, unitary PC inputs have highly variable amplitudes, with some large inputs that are 10 to 20 times the size of small inputs (**Fig. 1a-c, e-f,** n=83). This developmental transition indicates that PC-CbN synapses are refined during maturation, potentially because of plasticity mechanisms (Aizenman et al., 1998; Morishita and Sastry, 1996; Ouardouz and Sastry, 2000). The implications of such variability in the strength of PC-CbN synapses in cerebellar computation can be shown by a recent study describing how the cerebellum encodes vestibular and neck proprioceptive information (Zobeiri and Cullen, 2022). Each PC encode both vestibular and proprioceptive information, but some CbN neurons exclusively encode body motion and others encode vestibular information (Brooks and Cullen, 2009; Zobeiri and Cullen, 2022). It was proposed that this arises from the linear summation of converging PCs with different weights (Zobeiri and Cullen, 2022). Our observation of the variable distribution of PC-CbN inputs provide evidence for the requisite different weights of PC inputs.

The developmental changes in synaptic strengths we observe is reminiscent of other synapses. For retinal ganglion cell inputs to lateral geniculate nucleus, in young mice, numerous weak inputs innervate each LGN neuron, but in mature mice, this gives way to highly skewed input sizes that include some very strong inputs (Hooks and Chen, 2020; Jiang et al., 2022; Liang and Chen, 2020; Litvina and Chen, 2017). Excitatory inputs in the superior colliculus are refined in a comparable manner (Lu and Constantine-Paton, 2004). For inhibitory inputs to LSO neurons, in P1-5 mice there are many small and no large inputs, but by P9-14 the input sizes are highly skewed, and some inputs are very large (Gjoni et al., 2018; Kim and Kandler, 2003). In the cortex, input sizes are highly variable, and are often approximated by a lognormal distribution (Buzsáki and Mizuseki, 2014; Dorkenwald et al., 2022; Melander et al., 2021). Thus, the distribution of PC to CbN input sizes we observe in P23-32 mice, is similar in many ways to the distribution of input sizes of other types of synapses.

### Nonuniform PC input sizes elevates the basal firing rate of CbN neurons

An important consequence of having highly variable PC-CbN inputs is that it elevates the basal firing rates of CbN neurons (**Fig. 4 and Fig. 6**). With the same total inhibition, uniform small PC inputs are particularly effective at suppressing the firing of CbN neurons. Nonuniform sized PC inputs elevate the CV of inhibition, increasing the fluctuations of membrane potential and elevating the firing of CbN neurons. Simulations based on an integrate-and-fire model with a passive cell and a well-defined firing threshold were able to replicate the findings of dynamic clamp experiments. This suggests that CbN neuron firing is effectively driven by the membrane potential fluctuations generated by PC inputs, despite the presence of many excitatory inputs and intrinsic conductances. The high basal firing rate of CbN neurons generated by nonuniform sized inputs helps to explain the relatively high firing rates of CbN neurons *in vivo* amid strong PC inhibition (10-50 Hz) (Eccles et al., 1974; LeDoux et al., 1998; McDevitt et al., 1987; Rowland and Jaeger, 2008, 2005; Thach, 1975, 1970b, 1968). Membrane potential fluctuations have also been shown to be important to spike generation in other cell types (Kuhn et al., 2004; Tiesinga et al., 2000; Yarom and Hounsgaard, 2011).

### Implications of nonuniform input sizes on rate codes

Our findings established that there is an inverse linear relationship between the firing rates of each PC input and the targeted CbN neuron (**Fig. 5**) (Person and Raman, 2012b; Wu and Raman, 2017). The extent to which the firing rate of an individual PC input regulates the firing rate of a CbN neuron simply depends on the size of the input, and our simulations suggest that single PCs are capable of regulating the firing of a CbN neuron from 35 to 125 spikes/s (**Fig. 5g**; for a 37 nS input varied from 0 to 160 spikes/s). Regardless of the sizes of the input that are varied, the changes in the firing rate of CbN neurons can be readily explained by the alterations in the total inhibitory conductance. It was not obvious prior to these experiments that there would be such a simple relationship between input size, input firing rate, and output firing rate, given that CbN firing rates depend on both the amplitude and CV of inhibition. In practice, fluctuations in the inhibitory conductance are quite large for variable size inputs, and changing the rate of one input has a relatively small influence on the CV of the inhibition, so the magnitude of the inhibitory conductance dominates the effect. This model allows all different sized PC inputs to reliably convey a simple rate code, with the large PC inputs being particularly effective.

### Implications of nonuniform inputs on temporal codes

Previously it was also thought that an individual PC input has a small influence on the spike timing of a CbN neuron, and that it is necessary to synchronize the firing of many PCs to precisely entrain CbN firing (Heck et al., 2013; Person and Raman, 2012a). Our cross-correlation analysis shows that this is not the case: large inputs (30 nS) eliminated CbN neuron firing for several milliseconds, and even small inputs (3 nS) transiently reduced CbN neuron firing by almost 40% (**Fig. 2**). The sizes of PC inputs, the rapid kinetics of PC-CbN IPSCs (Person and Raman, 2012a), the high firing rate of PCs (Thach, 1968; Zhou et al., 2014), and the strong tendency of CbN neurons to depolarize (Raman et al., 2000) together shape the pattern of cross-correlograms and contribute to precise control of CbN neuron spike timing. Thus, PC-DCN synapses are well suited to regulating both the rate and precise timing of CbN firing.

Surprisingly, there was an excitatory response in CbN neuron firing preceding the suppression by PCs in the cross-correlogram of PC inputs and CbN neuron firing (**Fig. 2**). This excitation is a consequence of the autocorrelation function in PC firing (Ostojic et al., 2009), and it will be most prominent for rapidly firing cells (**Fig. 3**). When PCs fire at frequencies lower than 50 spikes/s, the excitatory component is small and lasts for several milliseconds, but when PCs fire at over 100 spikes/s, the excitatory component increases in amplitude and decreases in duration. This indicates the firing rates of PCs will affect the way they control the timing of CbN neuron firing. The observation that PCs in zebrin-regions of the cerebellum tend to fire faster than PCs in zebrin+ regions suggest different timing control of CbN neurons by zebrin+ and zebrin-PCs (Zhou et al., 2014). The effective excitation might also be present for other inhibitory synapses (Arlt and Häusser, 2020; Blot et al., 2016).

The observed distribution of PC to CbN synapse amplitudes, combined with the firing properties of PCs *in vivo*, allow us to make several predictions about PC-CbN neuron cross-correlograms *in vivo*. It is expected that PCs to CbN synapses are sufficiently large that prominent suppression should be apparent in cross-correlograms *in vivo*. The variability of PC-CbN synapses amplitudes suggests that there will be considerable variability in the extent and duration of spike suppression that is comparable to **Fig. 2**. Single large inputs are expected to eliminate CbN neuron firing for several milliseconds, but smaller, weaker connections are expected to be more prevalent. Our findings also predict a disinhibitory component preceding the suppression that will be particularly large for rapidly firing PCs.

### Implications for the influence of PC synchrony

Variable PC input sizes have important implications for how PC synchrony regulates the firing of CbN neurons. Firstly, the relative influence of PC synchrony on the firing rate of CbN neurons is reduced because variable input sizes elevate the basal firing rates of CbN neurons (**Fig. 4**, **Fig. 6**). Secondly, the effect of PC synchrony depends upon which inputs are synchronized. Synchronizing 50% of the small inputs (corresponding to 31% of the total inputs) increased CbN firing by just 13%, whereas synchronizing the two largest PC inputs to a CbN neuron increased the CbN firing rate by 21% (**Fig. 6k and 6l**). These findings suggest that a high degree of synchrony is not prerequisite for an appreciable influence. Therefore, studies that fail to detect a high degree of synchrony in PC simple spike firing *in vivo* (Herzfeld et al., 2023) do not exclude the physiological relevance of PC synchrony in regulating CbN neurons. Understanding how weighted synaptic strength of PC-CbN connections are distributed and their potentials to synchronize are crucial to probe the physiological relevance of PC synchrony in cerebellar computation.

## Methods

| Key Resources Table |  |  |  |  |
| --- | --- | --- | --- | --- |
| Reagent type (species) or resource | Designation | Source or reference | Identifiers | Additional information |
| Strain, strain background (M. musculus) | C57/BL6 | Charles River Laboratories | N/A |  |
| Chemical compound, drug | NBQX disodium salt | Abcam | Ab120046 |  |
| Chemical compound, drug | (R,S)-CPP | Abcam | Ab120160 |  |
| Chemical compound, drug | Strychnine hydrochloride | Abcam | Ab120416 |  |
| Chemical compound, drug | SR95531 (gabazine) | Abcam | Ab120042 |  |
| Chemical compound, drug | CGP 55845 hydrochloride | Abcam | Ab120337 |  |
| Chemical compound, drug | QX-314 chloride | Abcam | Ab120118 |  |
| Software, algorithm | IgorPro | Wavemetrics (https://www.wavemetrics.com/order/order_igordownloads6.htm) | RRID:SCR_000325 | Version 6.37 |

### Ethics

All animal procedures were carried out in accordance with the NIH and Animal Care and Use Committee (IACUC) guidelines and protocols approved by the Harvard Medical Area Standing Committee on Animals.

### Animals

C57/BL6 wild-type mice (Charles River Laboratories) of both sexes aged P10 to P32 were used for acute slice experiments (for dynamic clamp experiments, P26 to P32). All animal procedures were conducted in accordance with the National Institutes of Health and Animal Care and Use Committee guidelines and protocols approved by the Harvard Medical Area Standing Committee on Animals (animal protocol #1493).

### Slice preparation

Mice were anesthetized with ketamine / xylazine / acepromazine and transcardially perfused with warm choline ACSF solution (34°C) containing in mM: 110 Choline Cl, 2.5 KCl, 1.25 NaH_2_PO4, 25 NaHCO_3_, 25 glucose, 0.5 CaCl_2_, 7 MgCl_2_, 3.1 Na Pyruvate, 11.6 Na Ascorbate, 0.002 (R,S)-CPP, 0.005 NBQX, oxygenated with 95% O_2_ / 5% CO_2_. To prepare sagittal cerebellar nuclei slices, the hindbrain was removed, a cut was made down the midline of the cerebellum and brainstem, and the halves of the cerebellum were glued down to the slicing chamber. Sagittal slices (170 μm) were cut using a Leica 1200S vibratome in warm choline ACSF (34°C). Slices were transferred to a holding chamber with warm ACSF solution (34°C) containing in mM: 127 NaCl, 2.5 KCl, 1.25 NaH_2_PO_4_, 25 NaHCO_3_, 25 glucose, 1.5 CaCl_2_, 1 MgCl_2_ and were recovered at 34°C for 10 minutes before being moved to room temperature for another 20-30 mins until recordings begin.

### Electrophysiology

Whole-cell voltage/current-clamp recordings were performed on large neurons (>70 pF) in the lateral and interposed deep cerebellar nuclei. These large cells are primarily glutamatergic projection neurons (Baumel et al., 2009; Turecek et al., 2016; Uusisaari et al., 2007).

For voltage-clamp recordings of unitary PCs inputs to CbN neurons, Borosilicate glass electrodes (1-2 MΩ) were filled with a high chloride (E_Cl_ = 0 mV) internal containing in mM: 110 CsCl, 10 HEPES, 10 TEA-Cl, 1 MgCl_2_, 4 CaCl_2_, 5 EGTA, 20 Cs-BAPTA, 2 QX314, and 0.2 D600, adjusted to pH 7.3 with CsOH. BAPTA was included to prevent long-term plasticity (Ouardouz and Sastry, 2000; Pugh and Raman, 2006; Zhang and Linden, 2006). The osmolarity of internal solution was adjusted to 290-300 mOsm. Series resistance was compensated up to 80%, and the calculated liquid junction potentials were around 5 mV and were left unsubtracted. CbN neurons were held at −30 to −40 mV. All experiments were performed at 34-35°C in the presence of 5 μM NBQX to block AMPARs, 2.5 μM (R,S)-CPP to block NMDARs, 1 μM strychnine to block glycine receptors, and 1 μM CGP 55845 to block GABA_B_Rs, with a flow rate of 3-5 ml/min. A glass monopolar stimulus electrode (2–3 MΩ) filled with ACSF was placed in the white matter between the CbN and the cerebellar cortex to activate PC axons. Minimal stimulation was used to determine the amplitudes of single PC-CbN inputs. The stimulus intensity was adjusted so that synaptic inputs were activated in approximately half the trials in a stochastic manner. The sample size was achieved by performing as many recordings as possible within a limited period of time.

For dynamic clamp experiments, Borosilicate glass electrodes (3-4 MΩ) were filled with an internal containing (in mM) 145 K-gluconate, 3 KCl, 5 HEPES, 5 HEPES-K, 0.5 EGTA, 3 Mg-ATP, 0.5 Na-GTP,5 phosphocreatine-tris2, and 5 phosphocreatine-Na2, adjusted to pH 7.2 with KOH. The osmolarity of internal solution was adjusted to 290-300 mOsm. Series resistances were less than 15 MΩ and were not compensated. Voltages were corrected for a liquid junction potential of 10 mV. Cells were held at −65 to −75 mV between trials. All experiments were performed at 34-35°C in the presence of 5 μM NBQX, 2.5 μM (R,S)-CPP, 5 μM SR 95531 (Gabazine) and 1 μM strychnine to block most synaptic transmission.

### Dynamic clamp experiments

The total inhibitory conductance (200 nS) from all converging PCs in each CbN neurons was based on previous estimation of 40 PCs with a size of 5 nS (after depression) (Person and Raman, 2012a). Uniform size inputs were studied in **Fig. 3**, **Fig. 4c-g**, and **Fig. 6**. For the uniform inputs with different firing frequencies used in Fig. 3, the number of the inputs were adjusted so that the total inhibitory conductances were the same for all groups (after depression, 12 x 20 nS at 49 Hz, 9 x 20 nS at 83 Hz, 6 x 20 nS at 122 Hz, and 9 x 20 nS for Poisson inputs). For the uniform inputs with different sizes used in **Fig. 4c-g**, the input sizes were varied from 2.5 nS to 40 nS (after depression), and the number of inputs was varied to maintain a total inhibitory conductance of 200 nS. 40 PC inputs with a size of 5 nS were used in Fig. 6.

Dynamic clamp experiments with different size inputs were performed in **Fig. 2 a-e**, **Fig. 4 h-i**, **Fig. 5**, **Fig. 6**. To generate different size inputs reflecting the distribution of the unitary PC-CbN input conductances measured in P23-32 animals, the amplitudes of the unitary conductances were corrected for depression (x 0.4) and the effects of high Cl-internal (scale-down by a factor of 2.3) (Bormann et al., 1987; Gjoni et al., 2018; Sakmann et al., 1983). In experiments where the effects on average firing frequency was determined (**Fig. 4hi**), input sizes were randomly drawn from the experimentally determined distributions of input sizes (**Fig. 2f**, *red*) until the total inhibitory conductance reached 200 nS. In experiments where the spike triggered averages were to be determined, we used a simplified distribution. In **Fig. 2a-e** and **Fig. 5** we approximated the distribution with small (16 × 3 nS), medium (10 × 10 nS) and large (2 × 30 nS) inputs. Approximating the small inputs with 16 inputs of the same size made it possible to determine the spike triggered average with much better signal to noise than if the inputs were different sizes. We took a similar approach in **Fig. 6**, but we adjusted the number and size of medium size inputs to make it easier to assess the effects of 50% and 25% synchrony (16 × 3 nS small, 8 × 12 nS medium and 2 × 30 nS large inputs).

We based the timing of PC firing on *in vivo* recordings (**Fig. 2—figure supplement 1**). The average firing frequency ranges from 61 spikes/s to 180 spikes/s (**Fig. 2—figure supplement 1a-b**). The distribution of interspike intervals of firing (ISIs) in the 10 PCs were well approximated with lognormal functions (**Fig. 2—figure supplement 1c**), and the relationship between the mean and the standard deviation (sd) of lognormal distributions fits were well approximated with a linear function: sd = −0.00154 + 0.583*mean (**Fig. 2—figure supplement 1d**). Therefore, we used this linear function to calculate the σ of a desired firing frequency (1/mean) and generated an artificial ISI distribution of PC firing based on lognormal function with the designated mean and sd. The ISI distributions of PC firing used in **Fig. 2** and **Fig. 6** were artificial lognormal distributions with a firing frequency of 83 Hz (**Fig. 2—figure supplement 1c**) and 80 Hz (**Fig. 2—figure supplement 1e-f**), respectively. The three ISI distributions of PC firing with different firing frequencies used in **Fig. 3** are from *in vivo* recordings (**Fig. 2—figure supplement 1a-b**), and the Poisson distribution without a refractory period was generated with an exponential function aiming at a desired frequency (80 Hz). The ISI distribution of PC firing used in **Fig. 4** is from *in vivo* recordings with a firing frequency of 100 Hz (**Fig. 2—figure supplement 1a-b**). The ISI distributions of PC firing with different firing frequencies used in **Fig. 5** are artificial lognormal distributions generated from the μ and σ of desired frequencies. The approach is shown for artificial lognormal distributions of 40, 80, 120, and 160 spikes/s (**Fig. 2—figure supplement 1e-f**). Individual PC spike trains were generated by randomly drawing ISIs from the designated ISI distribution, and spike trains from each PC were combined as a final spike train with the inputs from all PCs. This spike train was then convolved with a unitary PC input with a rise time of 0.1 ms, a decay time of 2.5 ms (Khan et al., 2022). The reversal potential for inhibitory conductances was set at −75 mV (after correcting for junction potential).

Excitatory conductances were based on the AMPA component of mossy fiber (MF) EPSCs characterized in previous studies (Wu and Raman, 2017), with a rise time of 0.28 ms, a decay time of 1.06 ms, and an amplitude of 0.4 nS (reflecting depression). They estimated that 20-600 MFs converged on each CbN neuron, with unknown firing frequencies, so the excitatory conductances were relatively unconstrained. We adjusted the frequency of MF EPSCs so that the basal firing rate of CbN neurons was maintained at 20-40 Hz in the presence of the inhibitory conductance. The average baseline excitatory conductance was 20-30 nS.

To avoid a drastic increase in spike frequency and the following adaptation of CbN neurons resulting from big changes in the variability of conductances, we ramped up the CV of the inhibitory conductances in the beginning of each trial so that the firing rate of CbN increased gradually from the hyperpolarization state. Each conductance was repeated in the same neuron for 3-4 trials as technical replicates. For synchrony experiments, PC inputs were synchronized 100% in their spike times. Therefore, synchronizing 10 5 nS inputs is equivalent to having one big input with a size of 50 nS.

### Analysis

Recordings were obtained using Multiclamp 700B (Molecular Devices), sampled at 50 kHz and filtered at 4 kHz, and collected in Igor Pro (WaveMetrics). Dynamic-clamp recordings were performed with an ITC-18 computer interface controlled by mafPC in Igor Pro (WaveMetrics). Data were analyzed using custom-written scripts in MATLAB (MathWorks) and Igor Pro (WaveMetrics). Auto-correlation and cross-correlation analysis were performed by generating accumulative histograms of spikes distribution within a 20 ms time window centering all spikes from the reference file (self-reference for auto-correlation, and PCs spike times for cross-correlation), and normalized by the total spikes number of the reference file and the bin size of the histograms (that is, the Δt). All summary data are shown as the mean ± SEM unless otherwise indicated. The distributions of unitary PC input sizes in young and juvenile animals in **Fig. 1f** were compared with a Kolmogorov–Smirnov test. The unpaired t test was performed with Welch’s correction.

### Simulations

Simulations were performed (**Fig. 2f-h**, **Fig. 4j**, **Fig. 5g-h**, and **Fig. 6g-j, o-p)** to complement dynamic clamp experiments. A point-conductance, single-compartment model was generated to model the CbN neuron and its synaptic inputs. The membrane potential (*V*) of a CbN neuron with a membrane capacitance (*C*_*m*_) follows the equation:

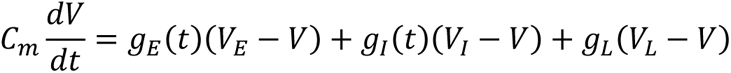

where *V*_*E*_ = 0 mV and *V*_*I*_ = −75 mV are the excitatory and inhibitory reversal potentials, and *g*_*E*_ (*t*) and *g*_*I*_(*t*) are time-dependent excitatory and inhibitory conductances. The leak was modelled by a constant leak conductance *g*_*L*_ with a reversal potential *V*_*L*_. A spike occurs when the membrane potential *V* of the model neuron reaches the threshold θ, at which point there is a refractory period of 2 ms during which the model neuron remains inactive, and the neuron is then reset to *V*_*r*_. *g*_*E*_ (*t*) and *g*_*I*_(*t*) were generated as described for dynamic clamp experiments. Simulations were performed using the BRIAN simulation environment in Python. Code can be accessed in GitHub: https://github.com/asemptote/PC-DCN-different-size-inputs.

In **Fig. 2f-h**, simulations were performed for 10 cells with input distributions drawn randomly from the list of empirical input sizes such that the total conductance was 200 nS. Each cell was run for 160,000 s and cross-correlograms were computed for each input. The parameters used were *C*_*m*_ = 50 pF, 20000 excitatory events per second, θ = −50 mV, *V*_*r*_ = −60 mV, *g*_*L*_ = 8.8 nS, and *V*_*L*_ = −40 mV. Inhibitory spike trains were randomly chosen by drawing ISIs from the empirically fitted lognormal distribution such that the mean rate was 83 Hz.

In **Fig. 4j**, simulations were performed for 100 cells with input distributions drawn randomly from the measured and corrected input sizes (**Fig. 2f**, *red*), along with 200 cells with varying numbers of uniform-size inputs such that the total conductance was 200 nS. Each cell was run for 10 s and the firing rates and inhibitory conductance CV were recorded. The generated conductance waves were saved for use in experiments in Fig. 4i. The parameters used were *C*_*m*_ = 200 pF, 23650 events per second, θ = −50 mV, *V*_*r*_ = −60 mV, *g*_*L*_ = 5 nS, and *V*_*L*_ = −10 mV. For the cells with varying input sizes, inhibitory spike trains were randomly chosen by drawing ISIs from the empirically fitted lognormal distribution such that the mean rate was 83 Hz, while for the cells with uniform-size inputs, the ISIs were drawn from an *in vivo* recording with a firing frequency of 100 Hz (**Fig. 2—figure supplement 1**).

In **Fig. 5g-h**, simulations were performed for 10 cells with input distributions drawn randomly from the list of empirical input sizes such that the total conductance was 200 nS. For each input to each cell, 16 simulations were run for 100 s, and the rate of a given input was varied between 0 and 160 Hz while keeping the other inputs at a fixed rate (80 Hz). The average firing rate, the mean inhibitory conductance and its CV were recorded. The parameters used were as **in Fig. 4**.

In **Fig. 6**, simulations were performed for 10 cells with input distributions drawn randomly from the list of empirical input sizes such that the total conductance was 200 nS. For each cell, 100 simulations were run for 1600 s whereby a random subset of the input sizes was synchronized, and the cross-correlation statistics were obtained as in **Fig.2**. 40 additional simulations were run in this manner for a cell with 40 uniform-size inputs, synchronizing a different number of inputs in each simulation. The parameters used were as in **Fig. 4**.

## Acknowledgements

We thank members of the Regehr lab, Indira Raman and Nicolas Brunel for comments on the manuscript. We thank Josef Turecek and Skyler Jackman for their initial work on charactering the unitary PC inputs to CbN neurons. This work was supported by grants from the NIH (R01NS032405 and R35NS097284 to W.G.R.).

## Author contribution

**Shuting Wu**: Conceptualization, Methodology, Formal analysis, Investigation, Writing - Original Draft, Visualization.

**Asem Wardak**: Methodology, Software, Formal analysis, Investigation, Visualization, Writing - review and editing.

**Mehak M. Khan**, Investigation, Writing - review and editing.

**Christopher H. Chen**: Investigation, Writing - review and editing.

**Wade G. Regehr**: Conceptualization, Resources, Funding acquisition, Project administration, Visualization, Writing - original draft, Writing - review and editing.

## Competing interests

The authors declare no competing interests.

## Notes

### Competing Interest Statement

The authors have declared no competing interest.

